# Targeting Apollo-NADP^+^ to image NADPH generation in pancreatic beta-cell organelles

**DOI:** 10.1101/2022.04.27.489691

**Authors:** Huntley H. Chang, Alexander M. Bennett, William D. Cameron, Eric Floro, Aaron Au, Christopher M. McFaul, Christopher M. Yip, Jonathan V. Rocheleau

## Abstract

NADPH/NADP^+^ redox state supports numerous reactions related to cell growth and survival; yet the full impact is difficult to appreciate due to organelle compartmentalization of NADPH and NADP^+^. To study glucose-stimulated NADPH production in pancreatic beta-cell organelles, we targeted the Apollo-NADP^+^ sensor by first selecting the most pH stable version of the single colour sensor. We subsequently targeted mTurquoise2-Apollo-NADP^+^ to various organelles and confirmed activity in the cytoplasm, mitochondrial matrix, nucleus, and peroxisome. Finally, we measured the glucose- and glutamine-stimulated NADPH responses by single and dual colour imaging of the targeted sensors. Overall, we developed multiple organelle-targeted Apollo-NADP^+^ sensors to reveal the prominent role of beta-cell mitochondria in determining NADPH production in the cytoplasm, nucleus, and peroxisome.

---

NADPH/NADP^+^ redox state provides electrons to biosynthetic reactions required for cell growth and proliferation, and to recharging antioxidants required for survival^1^. NADPH/NADP^+^ redox state is thought to be compartmentalized due to a lack of known NADPH or NADP^+^ membrane transporters, and the existence of organelle-specific enzymes for NADP^+^ biosynthesis and reduction^1,2^. Significant compartmentalization could create disparate impacts on organelle-specific anabolism and antioxidation. Our goal was to target a sensor we previously developed, Apollo-NADP^+^, to organelles to measure organelle-specific NADPH compartmentalization.

Pancreatic beta-cells metabolically respond to glucose to stimulate insulin secretion. This response includes a rise in NADPH/NADP^+^ ratio. Pancreatic beta-cells show limited G6PD activity^3^ and thus rely on cytoplasmic malic enzyme (ME)^4–6^ and/or isocitrate dehydrogenase (IDH1)^7^ activity driven by metabolite efflux from the mitochondria. Mitochondria generate their own NADPH using NADP^+^-dependent isocitrate dehydrogenase (IDH2) and nicotinamide nucleotide transhydrogenase (NNT)^8^ with previous work suggesting NNT dominates the glucose-stimulated response^9–12^. Thus, we aimed to measure the metabolic routes of beta-cell glucose-stimulated NADPH production and the extent of redox crosstalk between organelles.

Two genetically encoded NADH/NADP^+^ redox state sensors are available including the NADPH-sensitive iNAP sensor^13,14^ and the NADP^+^-sensitive Apollo-NADP^+^ sensors^15^. Apollo-NADP^+^ sensors are based on NADP^+^-induced dimerization of enzymatically inactive glucose-6-phosphate dehydrogenase (G6PD). Tagged with a single fluorescent protein, Apollo-NADP^+^ responds through changes in steady-state fluorescence anisotropy due to FRET between homologous fluorescent proteins (homoFRET). Importantly, anisotropy sensors consume less spectral bandwidth than conventional ratiometric sensors and can be swapped to distinct colors (i.e., they are spectrally tuneable), both of which facilitate multiparameter imaging^15–17^. In this study, we target Apollo-NADP^+^ to various organelle compartments. We subsequently use simultaneous two-colour imaging to show a dominant role of mitochondrial NADPH generation in setting the glucose-stimulated responses in the cytoplasm, nucleus, and peroxisome.

## MATERIALS AND METHODS

### Molecular biology of sensor targeting

We cloned Cerulean and mVenus-tagged Apollo-NADP^+^ into the *NheI/XhoI* sites of pcDNA3.1. The mTurqoise2 and Cerulean3 versions were subsequently created by exchanging the fluorescent protein using the *AgeI/Xho1* sites. Organelle targeting sequences were introduced into the mTurquoise2 version of Apollo-NADP^+^. Smaller tags were introduced using cloning primers, including: the nuclear targeting sequence of SV40 T-antigen ‘PKKKRK’ into the N-terminus^18^, the peroxisome targeting sequence – SKL into the C-terminus^19^, and the ER retention KDEL sequence into the C-terminus. Larger tags were cloned into the *NheI* and *BamHI/HindIII* sites, on the N-terminus of Apollo-NADP^+^. We cloned in the mitochondrial matrix targeting sequence from pKillerRed-dMito (a gift from Peter Kim, SickKids Hospital, Toronto) to get four copies of the mitochondrial targeting sequence. The membrane targeting sequence of neuromodulin was cloned from pEYFP-Mem, and the ER targeting sequence of calreticulin was cloned from pECFP-ER (Clonetech, San Jose, CA). All constructs were confirmed by sequencing. All plasmid amplification was performed using 5-alpha competent E. coli (NEB, Ipswich, MA, C2987I)

### Cells and cell culture

INS-1E (obtained from lab of Michael B. Wheeler, RRID: CVCL_0351) and AD293 (Agilent, Santa Clara, CA, #240085) cells were cultured in RPMI and DMEM media, respectively under humidified 5% CO_2_ at 37°C. For imaging, cells were plated into No 1.5 glass bottom dishes (MatTek Corporation, Ashland, MA) and transfected the next day (0.5-1.0 μg/dish/plasmid) using Poly-Jet reagent (for AD293s) (SignaGen, Frederick, MD) or Lipofectamine 3000 (for INS-1Es) (Invitrogen, Burlington, Canada) and incubated for 24 hr. The media was replaced with fresh media and the cells were left to recover for a further 24 hr prior to imaging.

### Clamping the intracellular pH of live cells

Cytoplasmic pH was assessed using the Intracellular pH Calibration Buffer Kit (Invitrogen, Burlington, Canada). Cytoplasmic pH was varied as described previously^20^. Cell loading solutions were prepared to the specified pH using 10 μM valinomycin and 10 μM nigericin and the buffer solutions provided in the kit. Cells were incubated in the specified pH solutions for 5 min to allow for equilibration of intracellular and extracellular pH levels before imaging.

### Islet isolation, dispersion, and transduction

Animal procedures were approved by the Animal Care Committee of the University Health Network, Toronto ON, Canada, in accordance with the policies of the Canadian Council on Animal Care (Animal Use Protocol #1531). Pancreatic islets were isolated from 8 to 12-week-old C57Bl/6J (Jackson Lab, Bar Harbor, ME, RRID: IMSR_JAX:000664) male mice by collagenase digestion (Roche Applied Science)^21^. Isolated islets were transferred into an Eppendorf tube (1.5 ml) containing 50 μl RPMI islet media and 50 μl Trypsin/EDTA, immersed in a water bath at 37 °C, and shaken gently by hand for 12 min. The dissociated islet slurry was topped up to 1 ml total volume with RPMI islet media. Glass bottom plates (48-well) were coated with Poly-D-Lysine hydrobromide (Sigma-Aldrich, Product #P6407) for 1 h at 37°C to promote cell adhesion. The dispersed cells were transduced with adenovirus (24 hr at 1:10 dilution or 2×10^7^ IFU/ml)^22^. Viral titers were measured using the Adeno-X rapid titer kit (Clontech) following manufacturer protocol.

### Cellular imaging

All imaging was done in BMHH imaging buffer (125 mM NaCl, 5.7 mM KCl, 2.5 mM CaCl_2_, 1.2 mM MgCl_2_, 0.1% BSA, and 10 mM HEPES at pH 7.4). Single colour two-photon fluorescence anisotropy imaging was performed using an LSM710 confocal microscope (Zeiss, Toronto, Canada) equipped with a tunable Chameleon laser (Coherent, Santa Clara, CA). Images were collected using 840 or 950 nm excitation and a 63×/1.4 NA oil-immersion objective. Fluorescence was collected using the non-descanned binary GaAsP BiG (Zeiss, Toronto, Canada) detector with a custom-built filter cube containing an infrared light–blocked Cerulean (ET480/40M-2P) or Venus (ET525/50M-2P) emission bandpass filter (Chroma, Bellows Falls, VT), polarizing beam-splitter (Edmund Optics, Barrington, NJ) and clean-up polarizers (Chroma, Bellows Falls, VT). Simultaneous two-colour fluorescence anisotropy imaging was performed using a custom-built widefield RAMM microscope (ASI) equipped with excitation LEDs (405, 505, and 590nm), and excitation polarizing filter (Edmund Optics, Barrington, NJ). Images were collected using a 60×/1.42 NA oil-immersion or a 40×/0.75 NA air objective lens (Olympus, Richmond Hill, Canada). Fluorescence was passed through a Cerulean (ET470/24M) or Venus (ET535/30M) emission filter on a filter wheel, and subsequently split using an Optosplit II (Cairn, Faversham, UK) to simultaneously collect parallel and perpendicular emission light on separate regions of an IRIS 15 CMOS camera (Teledyne Photometrics, Tucson, AZ).

### Image analysis

Parallel (*I*_*‖*_) and perpendicular (*I*_*⊥*_) fluorescence intensity images were analyzed with a custom ImageJ plugin. These are available at https://github.com/RocheleauLab/Optosplit-Anisotropy-Analysis-scripts. The images were background corrected using a rolling ball filter. Pixel-by-pixel anisotropy (r) was calculated using the background corrected intensities: r = (I_‖_ – GI_⊥_)/(I _‖_ + 2G_⊥_)^23^. The G factor for the two-photon microscope was measured by exciting samples with both vertically (V) polarized and horizontally (H) polarized light and collecting the polarized emission^24,25^. The G-factor for the widefield microscope was calculated using fluorescein solutions, simplifying the standard anisotropy equation to G = *I*_*‖*_ / *I*_*⊥*_. For images collected using the 1.42 NA objective lens, we used additional correction factors (K_a_, K_b_, and K_c_) to account for the blurring of parallel and perpendicular intensities^23^. Regions of interest (ROI) were selected using the parallel intensity images to avoid selection bias.

### Statistics

All data are presented as mean ± S.E.M based on at least three separate experiment days (i.e., n=3 independent trials). Each independent trial consists of at least 5 to more than 30 cells (5-30 technical replicates). Iglewicz and Hoaglin’s robust test for multiple outliers was used to detect outliers from technical replicates. Statistical significance was determined using Prism 6 (GraphPad, San Diego, CA). Post-hoc tests were used to determine significance between groups following 1- or 2-way ANOVA. Tukey’s multiple comparison tests were used when comparing between all groups. Dunnett’s multiple comparison tests were used when comparing the control group vs other groups only. Sidak’s multiple comparison tests were used for repeated measures ANOVA. Anisotropy measurements were confirmed as normally distributed using a normal probability plot and D’Agostino-Pearson normality tests.

## RESULTS

### Determining pH sensitivity of Apollo-NADP^+^

Fluorescent proteins are variably sensitive to pH, which could result in potential artifacts in the application of genetically encoded sensors. To select a fluorescent protein for Apollo-NADP^+^ targeting, we first measured the impact of pH on the steady-state fluorescence anisotropy of fluorescent protein tandem-dimers (EGFP, mVenus, Cerulean, Cerulean3, and mTurquoise2) (**Fig. 1a-b**). These tandem-dimers are connected by a short linker to ensure homoFRET^26,27^, thus mimicking the low anisotropy of dimeric Apollo-NADP^+^ (**Fig. 1a**). These data show consistently low anisotropy in the blue tandem-dimers (Cer1, Cer3, mTurq2) across the full range of pH (**Fig. 1b**). In contrast, the EGFP and mVenus tandem dimers showed a progressive increase in anisotropy below pH 6.0. This is consistent with the lower pKa values of the bluer fluorescent proteins (3.7 to 4.7) compared to EGFP and mVenus (6.0 and 5.5, respectively). Dropping pH below the pKa results in a fractional decrease in molecular brightness (“darkening”) of the fluorophore and an artifactual rise in anisotropy.

**Figure 1.**
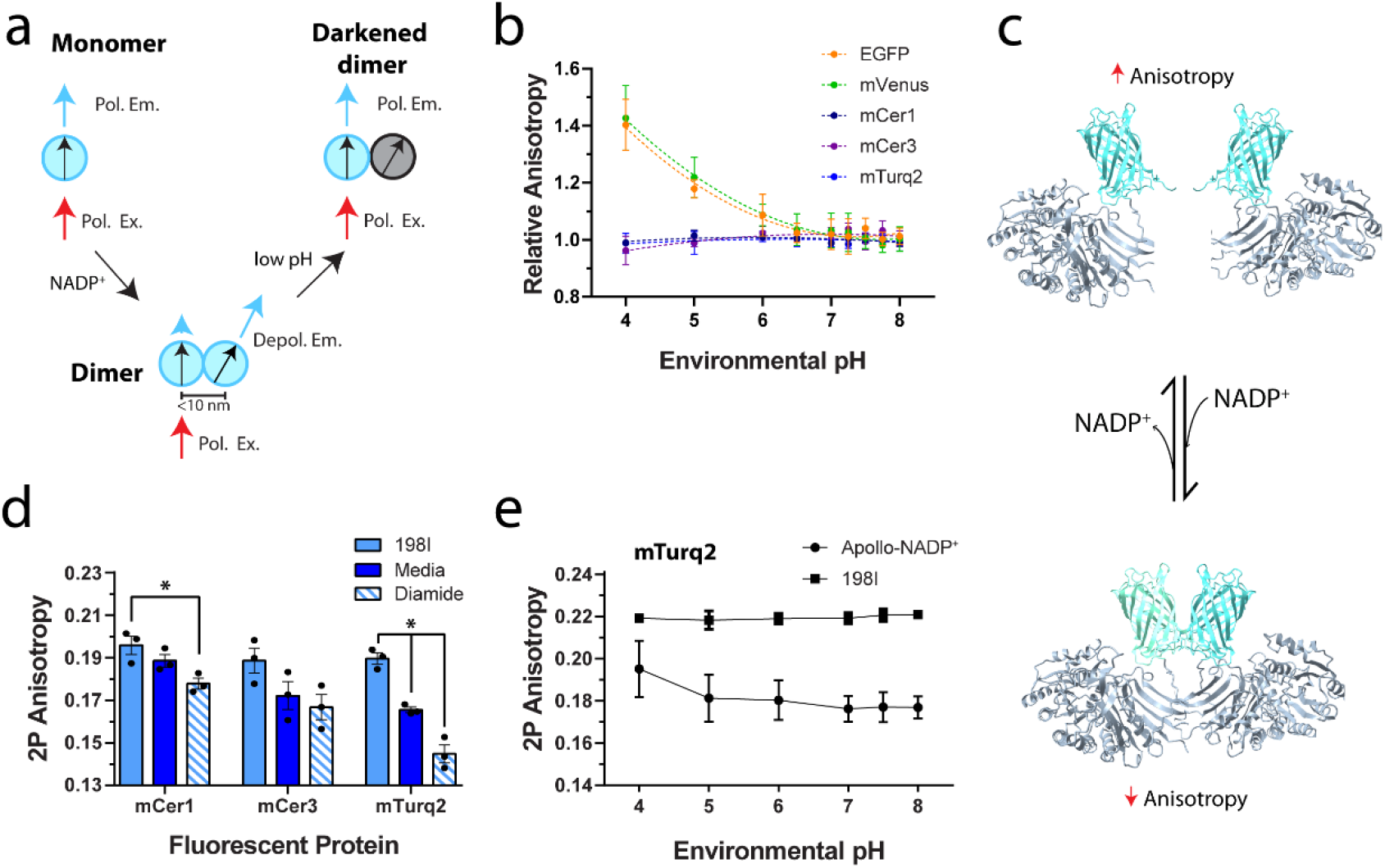
pH optimization of Apollo-NADP^+^. **(a)** Polarized excitation of fluorescent proteins results in polarized emission and high steady-state fluorescence anisotropy (Monomer). HomoFRET occurs when the proteins are within 10 nm causing depolarization of the emission and a drop in steady-state anisotropy (Dimer). A drop in pH below the pKa results in “darkening” (Darkened dimer) and an artifactual increase in anisotropy. **(b)** Tandem-dimers of EGFP, mVenus, mCerulean (mCer1), mCerulean3 (mCer3), and mTurquoise2 (mTurq2) were expressed in AD293 cells and imaged under varying pH. **(c)** The Apollo-NADP^+^ sensor, comprised of enzymatically inactivated human G6PD tagged with fluorescent protein, responds to NADP^+^ through allosteric dimerization. The crystal structures of human G6PD (PDB 6E07) and yellow fluorescent proteins (PDB 3V3D) are shown. **(d)** Apollo-NADP^+^ and monomeric R198P-G6PD (R198I) tagged with mCer1, mCer3 and mTurq2 were imaged in AD293 cells in media. Apollo-NADP^+^ was subsequently imaged after treatment with diamide (5 min, 5 mM), n = 3. **(e)** Cells expressing the mTurq2 versions of Apollo-NADP^+^ and R198I were exposed to 5mM diamide and imaged under varying pH, n = 3. The * denotes significance <0.05.

To further select from the blue fluorescent proteins, we compared the responses of Cer1-, Cer3-, and mTurq2-tagged Apollo-NADP^+^ in response to NADPH oxidation by diamide (**Fig. 1c-d**). NADP^+^-induced dimerization of Apollo-NADP^+^ decreases the steady-state fluorescence anisotropy due to homoFRET (**Fig. 1c**). The blue sensors all showed responses to diamide treatment compared to monomeric control (R198I) and sensor in normal media, with the largest responses found in cells expressing mTurq2-Apollo-NADP^+^ (**Fig. 1d**). These responses tracked well with the quantum yield (QY) of each fluorescent protein (mTurq2 (0.93) > mCer3 (0.87) >> mCer (0.49))^28^. This trend is consistent with the single fluorescent protein acting as both donor, impacting Förster distance (R_0_ ∝ (QY_D_)^½^), and acceptor, impacting the energy transmitted as depolarized fluorescence (I_A_ ∝ QY_A_). We subsequently settled upon the mTurq2 version as the most dynamic sensor.

We were concerned pH could also impact dimerization of the catalytically inactive G6PD. To determine the impact of pH on sensor dimerization, we next compared the fluorescence anisotropy of Apollo-NADP^+^ (mTurq2-Apollo-NADP^+^) and R198I in cells treated with diamide (**Fig. 1e**). These data show stable sensor dimerization across a wide range of pH, with increased anisotropy found below pH 5.0. This suggests G6PD, which translocates between the cytoplasm and peroxisome, evolved to withstand variations in pH^29–31^. Overall, mTurq2-Apollo-NADP^+^ was the most dynamic of the pH stable sensors, and thus was chosen for subsequent organelle targeting.

### Organelle targeting of mTurq2-Apollo-NADP^+^

We added targeting sequences for the mitochondrial matrix, nucleus, ER lumen, plasma membrane, and peroxisome to mTurq2-Apollo-NADP^+^. To confirm successful organelle targeting, we imaged each sensor in AD293s (**Fig. 2a**). These images show organelle-specific localization and morphology^32^. To validate the activity of the sensors we imaged in INS-1E beta-cells a corresponding set of targeted controls (blue) and sensor in response to glucose (0, 1, and 15 mM; red) and hydrogen peroxide (0.1, 10, and 1000 μM H_2_O_2_; yellow) **(Fig. 2b**). The controls included: a similarly targeted R198I monomer, the sensor in culture media, and diamide-induced sensor dimers. These controls defined the photophysical range of each sensor (i.e., monomer-to-dimer), except in the mitochondria and peroxisome where R198I did not consistently target the organelle. The inability of R198I to express in these organelles is consistent with an already destabilized construct being unable to fold in these organelles^33^. Notably, the anisotropies of the targeted R198I constructs were consistent across the other organelles (e.g., the cytoplasmic anisotropy was consistent with the nuclear anisotropy), and thus any of the other versions could serve as a monomeric control for the mitochondria and peroxisome.

**Figure 2.**
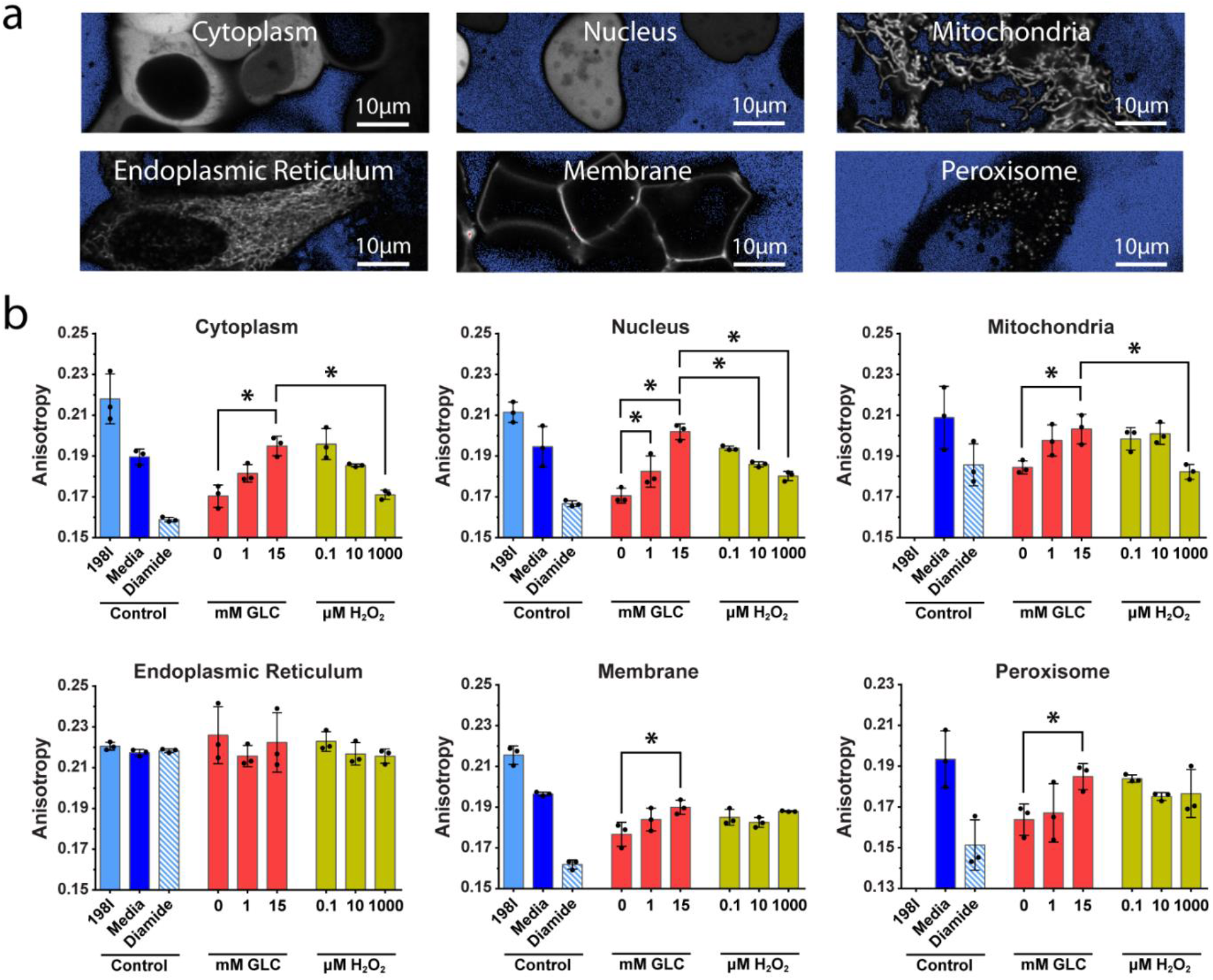
Characterization of organelle-targeted Apollo-NADP^+^ constructs. **(a)** Representative images of mTurq2-Apollo-NADP^+^ targeted to various subcellular compartments and expressed in AD293 cells. **(b)** mTurq2-Apollo-NADP^+^ and R198I constructs targeted to each organelle were expressed in INS-1E beta-cells. Controls included the R198I monomeric control (198I, where expressed), Apollo-NADP^+^ in full media (media), and 5 mM diamide (diamide). The sensor was imaged in sequential response to glucose and H_2_O_2_, n = 3. Not shown are the R198I values for the mitochondria (0.178±.002) and the peroxisome (0.097±.002) as the low intensity of the images and the variance in anisotropy suggested these constructs did not properly refold when targeted to these organelles. The * denotes significance <0.05.

Sensor targeted to the cytoplasm, mitochondria, and nucleus showed consistent rises in anisotropy induced by glucose and falls in anisotropy induced by H_2_O_2_ (**Fig. 2b, top**). While the nuclear and cytoplasmic responses mirrored each other, the mitochondrial response to H_2_O_2_ was dampened, only showing a significant response at 1000 μM H_2_O_2_. This could be due to dampening by cytoplasmic H_2_O_2_-scavenging prior to reaching the organelle and by greater antioxidant capacity at high glucose. In contrast, ER-targeted sensor showed an elevated anisotropy with no response to any of these treatments (**Fig. 2b, bottom left**). Consistent with non-functional sensor in the ER, INS-1E and AD293 cells also showed limited response to cortisone/hydrocortisone (data not shown), a treatment that affects ER NADPH/NADP^+^ redox state via 11β-HSD-1^34^. Membrane-targeted sensor showed a statistically significant response to glucose; however, the sensor did not subsequently respond to H_2_O_2_ suggesting an irreversible/slow recovery of monomerization due to molecular crowding in 2D at the membrane surface (**Fig. 2b, bottom middle**). Finally, the peroxisomal sensor showed a significant response to high glucose, but a much smaller response to H_2_O_2_. The glucose response is consistent with peroxisomes generating NADPH using G6PD and/or IDH1^29–31,36^. The smaller H_2_O_2_ response is consistent with catalase activity in peroxisomes detoxifying H_2_O_2_ without consuming NADPH. Overall, these data show functional Apollo-NADP^+^ sensors in the cytoplasm, nucleus, mitochondrial matrix, and peroxisome.

### Imaging organelle-specific Apollo-NADP^+^ to explore glucose-and glutamine-stimulated NADPH responses

Due to limited G6PD activity in beta-cells glucose-stimulated NADPH production primarily depends on mitochondrial efflux of citrate/isocitrate and IDH1 activity^3^. To measure this metabolic route, we co-expressed cytoplasmic mVenus-Apollo-NADP^+^ and mitochondrial mTurq2-Apollo-NADP^+^ in INS-1E cells and simultaneously imaged the glucose-stimulated temporal responses (**Fig. 3a and 3b**). These data show NADPH generation plateaus in the mitochondrial matrix (∼5 min, blue) prior to the cytoplasm (∼9 min, yellow). Both responses were significantly diminished by the mitochondrial pyruvate transport inhibitor UK5099. Together, these data are consistent with cytoplasmic NADPH production via mitochondrial metabolism of pyruvate.

**Figure 3.**
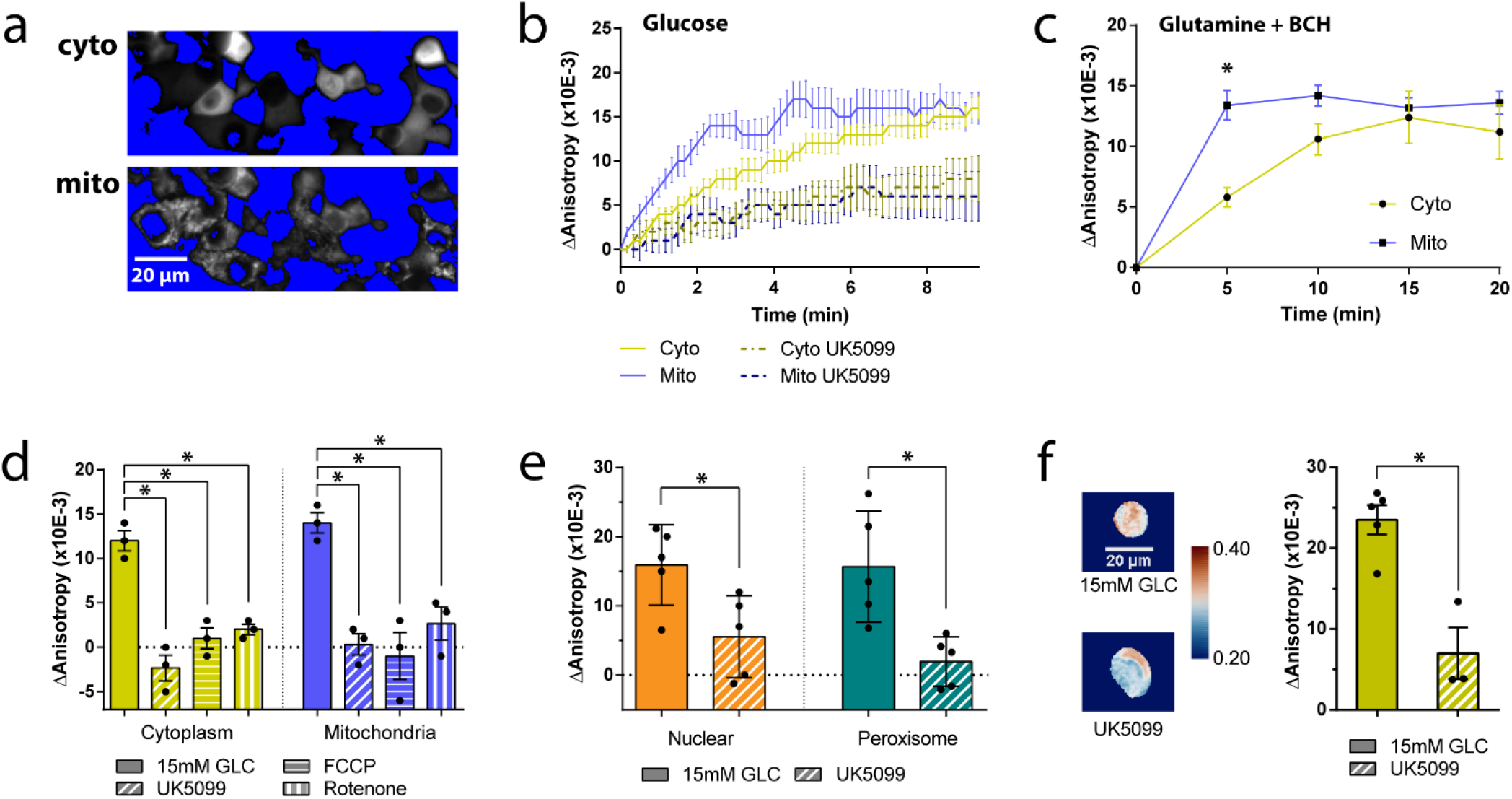
Using organelle-specific Apollo-NADP^+^ to determine the role of mitochondrial metabolism on NADPH responses. **(a)** Representative images of INS1E cells co-expressing mVenus-Apollo-NADP^+^ (Cyto) and mTurq2-Apollo-NADP^+^ (Mito) with similar thresholding and a 1 pixel median filter. **(b)** The temporal response to 15 mM glucose bolus was compared to cells pre-treated for 1 hr with 50 μM UK5099. Data are reported as mean change in anisotropy (ΔAnisotropy (×10^−3^)) ± S.E.M., n = 5. **(c)** Temporal response to 10 mM BCH in cells pre-treated with glutamine (5 mM, 1 hr), n = 3. **(d)** The anisotropy response to glucose (1 to15 mM, 5 min) in control cells and cells pre-treated with UK5099 (50 μM, 1 hr), FCCP (1 μM, 5 min), and rotenone (1 μM, 5 min), n = 3. **(e)** The anisotropy responses to a 10 (nuclear) or 20 (peroxisome) min glucose bolus in cells pretreated with UK5099, n = 5. **(f – left)** Representative anisotropy images of dispersed mouse islet cells transduced with cytoplasmic Apollo-NADP^+^ with similar thresholding and a 1 pixel median filter. Both cells were treated with 15 mM glucose (5 min) while the bottom cell was also pretreated with UK5099 (50 μM, 1 hr). Colour map is *vik*^35^. **(f – right)** Anisotropy responses to 15 mM glucose in controls or cells pre-treated with UK5099, n = 3-5. Control replicates are shared with **Fig. 4f** since the data was collected at the same time. The * denotes significance < 0.05.

Glutamine is a mitochondrial metabolite that enters the TCA cycle through glutamate dehydrogenase (GDH) activity when triggered by the non-metabolizable leucine analogue BCH^37^. To confirm mitochondrial metabolism leads to cytoplasmic NADPH generation, we measured the cytoplasmic and mitochondrial response to glutamine and BCH (**Fig. 3c**). These data show glutamine stimulates mitochondrial NADPH prior to the cytoplasm consistent with the cytoplasmic response depending on mitochondrial metabolism. To further explore the link between mitochondrial and cytoplasmic NADPH generation, we simultaneously imaged the cytoplasmic and mitochondrial responses in cells treated with UK5099 and various electron transport chain inhibitors (**Fig. 3d**). UK5099 abolished the cytoplasmic and mitochondrial responses. The glucose-stimulated cytoplasmic and mitochondrial responses were also both blocked by collapse of the mitochondrial membrane potential (FCCP) and inhibition of the electron transport chain (rotenone). Collectively, these data suggest that both mitochondrial membrane potential and mitochondrial NADPH accumulation are both required for a subsequent response in the cytoplasm. To measure the impact of pyruvate entry into mitochondrial metabolism on other organelles, we imaged glucose-stimulated responses in the nucleus and peroxisome in the presence of UK5099 (**Fig. 3e**). These data show blocking pyruvate entry into mitochondrial metabolism abolishes responses in the nucleus and peroxisome. Overall, these data suggest mitochondrial NADPH accumulation precedes cytoplasmic, nuclear, and peroxisomal NADPH generation consistent with dependence on mitochondrial metabolism (e.g., citrate/isocitrate efflux).

To validate our findings in primary tissue, we imaged the glucose-stimulated response of dispersed mouse islet cells transduced with cytoplasmic mVenus-Apollo-NADP^+^ (**Fig 3f**). As expected, the glucose-stimulated response was abolished by UK5099 consistent with cytoplasmic NADPH generation in primary beta-cells due to mitochondrial pyruvate metabolism.

### Imaging organelle specific Apollo-NADP^+^ to probe the impact of fatty acids on glucose-stimulated NADPH response

Fatty acids impact glucose-stimulated NADPH production through inhibition of pyruvate dehydrogenase (PDH) activity by acetyl-CoA^38^, and through inhibition of mitochondrial NNT activity at higher chain lengths^39^ (**Fig. 4a**). To measure the impact of PDH inactivation on glucose-stimulated NADPH responses, we co-expressed cytoplasmic (mVenus) and mitochondrial (mTurq2) sensors to simultaneously image the glucose-stimulated responses of octanoate treated cells (**Fig. 4b-c**). These data show octanoate decreases the glucose-stimulated cytoplasmic response with minimal impact on the mitochondrial response. Consistent with PDH inactivation, DCA restored the cytoplasmic response while leaving the mitochondrial response unaffected. The inability of octanoate to block mitochondrial NADPH generation suggested a prominent role for NNT. NNT converts NADH to NADPH using mitochondrial membrane potential, thus linking mitochondrial energetics to antioxidant capacity^8^. To confirm the role of NNT in the glucose-stimulated mitochondrial NADPH response, we next pretreated cells with palmitate, a longer chain fatty acid well-established to inhibit NNT activity (**Fig. 4d-e**)^39^. Palmitate requires transport by carnitine palmitoyltransferases, resulting in a longer pre-incubation time. Palmitate abolished both the cytoplasmic and mitochondrial glucose-stimulated responses consistent with inhibition of PDH and NNT (via palmitoyl-CoA). In contrast to octanoate, the cytoplasmic response in the presence of palmitate was unaffected by DCA. We therefore postulate that the cytoplasmic response depends on both entry of pyruvate into the TCA cycle and mitochondrial NADPH generation by NNT.

**Figure 4.**
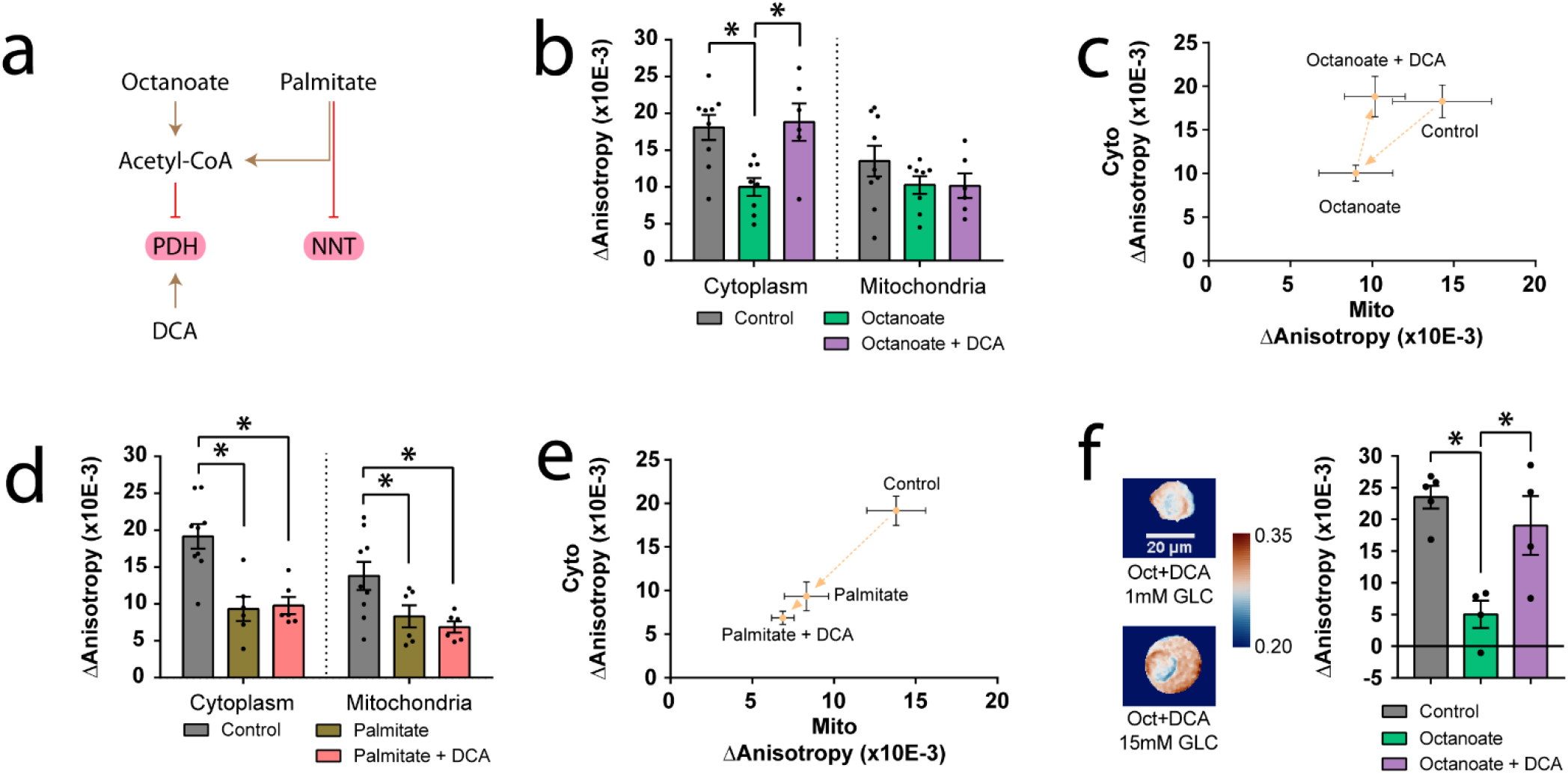
Using organelle-specific Apollo-NADP^+^ to determine the roles of non-anaplerotic fatty acids and NNT activity on cytoplasmic and mitochondrial NADPH. **(a)** Acetyl-CoA produced by beta-oxidation of both fatty acids will inhibit PDH activity with DCA reversing this effect. Longer chain fatty acid palmitate is also a direct inhibitor of NNT. **(b)** Simultaneous imaging of cytoplasmic and mitochondrial anisotropy responses to 15 mM glucose (15 min) in cells pre-treated with octanoate (400 μM, 30 min), octanoate and 1 μM DCA (30min), or no treatment (control). Data are reported as the mean change in anisotropy (ΔAnisotropy (×10^−3^)) ± S.E.M., n = 6-9. **(c)** The same anisotropy changes from (b) shown in scatter plot. Dotted arrows denote the order they are described in the results. **(d - e)** Simultaneous imaging of cytoplasmic and mitochondrial anisotropy responses to 15 mM glucose (15 min) in cells pre-treated with palmitate only (400 μM, 4 hr), or palmitate (4 hr) followed by 1 μM DCA (30 min) after a 15mM glucose bolus, n = 5-9. **(f – left)** Representative anisotropy images of dispersed mouse islets transduced with cytoplasmic Apollo-NADP^+^ with similar thresholding and a 1 pixel median filter. Cells were stimulated with 15 mM glucose (15 min) after pre-treatment in 400 μM octanoate (30min, top) or 400 μM octanoate + 1 μM DCA (30min, bottom). **(f – right)** Anisotropy responses of controls compared to cells pre-treated with octanoate or octanoate + DCA, n = 4-5. Control replicates are shared with **Fig. 3f** since the data was collected at the same time. The * denotes significance < 0.05.

To validate these findings in primary tissue, we imaged the impact of octanoate on the glucose-stimulated response of dispersed mouse islet cells transduced with cytoplasmic mVenus-Apollo-NADP^+^ (**Fig. 4f**). Octanoate significantly diminished the glucose-stimulated NADPH response, and this response was restored by activation of PDH using DCA. These data are again consistent with the cytoplasmic NADPH response being dependent on mitochondrial efflux in primary cells.

## DISCUSSION AND CONCLUSION

Our goal was to target Apollo-NADP^+^ to organelles to measure NADPH generation and compartmentalization in beta-cells. In contrast to the cytoplasm^40^, ER^41^ and nucleus^42^, the mitochondrial matrix^43^ and peroxisome^44^ can show significant variability in physiological pH. This variability complicates the application of genetically encoded sensors in these organelles. Although some pH sensitivity is not insurmountable, the methods to correct for it (e.g., co-expressing a pH sensor or an inactivated sensor)^14,40^ work against the ability to track single cells and perform multiparametric imaging. Since Apollo-NADP^+^ sensors are single colour senors, we were able to overcome this limitation by swapping in different fluorescent proteins. In general, the bluer fluorescent protein dimers (mCer, mCer3, and mTurq2) were more pH-stable than the greener fluorescent protein dimers (mVen, EGFP) likely due to lower pKa (pKa∼3.2)^28,45^. We subsequently settled upon the mTurq2 version as the most dynamic sensor noting the sensor responses tracked well with the QY of each fluorescent protein (mTurq2 (0.93) > mCer3 (0.87) >> mCer (0.49))^28^. This trend is consistent with the single fluorescent protein acting as both donor to impact Förster distance (R_0_ ∝ (QY_D_)^½^) and acceptor to impact the energy transmitted as depolarized fluorescence (I_A_ ∝ QY_A_). Thus, we targeted the mTurq2 version of the sensor due to having the highest pH stability and apparent dynamic range.

Highlighting the versatility of Apollo sensors, we proceeded with single (mTurq2) and dual colour (mTurq2 and mVenus) imaging of organelle-targeted Apollo-NADP^+^ sensors to examine beta-cell NADPH production. We showed glucose- and glutamine-stimulated NADPH production in the mitochondria prior to the cytoplasm, with both the glucose-stimulated responses blocked by the pyruvate transport inhibitor UK5099. These data are fully consistent with low G6PD activity in beta-cells leaving glucose-stimulated NADPH production in the cytoplasm dependent on mitochondrial efflux. Pyruvate enters beta-cell mitochondrial refilling of TCA cycle intermediates (i.e., anaplerosis), triggering mitochondrial citrate/isocitrate efflux and subsequent production of cytoplasmic NADPH via ME and/or IDH1. To examine the source of glucose-stimulated mitochondrial NADPH generation, we measured the impact of short (octanoate) and longer chain (palmitate) fatty acids on the glucose-stimulated mitochondrial and cytoplasmic responses. We postulated that inhibition of PDH by acetyl-CoA would lessen cytoplasmic NADPH response by decreasing metabolic flux through citrate synthase and/or increasing isocitrate consumption by NAD^+^-dependent isocitrate dehydrogenase (IDH3). Consistently, blocking PDH through the short chain fatty acid octanoate inhibited cytoplasmic, but not mitochondrial, NADPH generation. Additionally, DCA treatment led to PDH activation and rescued the cytoplasmic response. Finally, previous work suggests that NNT dominates glucose-stimulated mitochondrial NADPH production^6,8,46^. Consistently, the glucose-stimulated cytoplasmic and mitochondrial NADPH responses were abolished by inhibiting NNT using palmitate and by blocking mitochondrial membrane potential.

The glucose-stimulated nuclear and peroxisomal responses were also blocked by UK5099. Notably, the nuclear response mirrored the cytoplasmic response consistent with previous work suggesting NADPH is readily diffusible between cytoplasmic and nuclear compartments^14^. Thus, the nuclear targeted sensor could potentially serve as a useful surrogate for the cytoplasmic sensor to improve spatial separation during multiparameter imaging and to better isolate individual cell responses in tissues. However, substitution of the nuclear sensor will need to be fully considered in the context of varied compartmentalization of glutathione in the nucleus during the cell cycle^13,47,48^. Overall, this work expands the Apollo-NADP^+^ family of sensors while suggesting a dominant role of mitochondrial metabolism in setting beta-cell NADPH/NADP^+^ redox state across organelles.

## AUTHOR INFORMATION

### Author Contributions

#H.H.C and A.M.B. contributed equally to this work. H.H.C., A.M.B., W.D.C., and J.V.R. designed research; H.H.C., A.M.B., W.D.C., and E.F. performed research; H.H.C., A.M.B., W.D.C., and E.F. analyzed data; A.A., C.M.M., and C.M.Y provided technical expertise, and H.H.C. and J.V.R. wrote the paper.

### Notes

The authors declare no competing interests.

## ACKNOWLEDGEMENTS

These studies were supported by NSERC Discovery Grants to JVR (RGPIN-2016-371705) and CMY (RGPIN-2015-043), by a CIHR project grant to JVR (DOL 409157) and by a NSERC Research Tools and Instrumentation Grant (2018-00846) to JVR. We thank Yufeng Wang and Vidhant Pal for assistance with islet isolation expertise.

## FOR TABLE OF CONTENTS ONLY

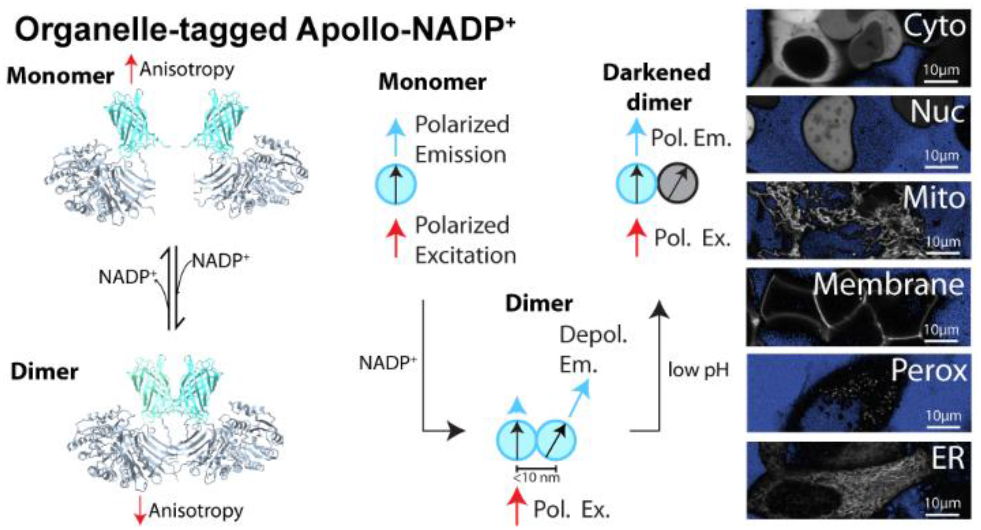

